# Differential multi-omics analysis of pulmonary arterial hypertension microvascular endothelial cells for differential drug response

**DOI:** 10.64898/2026.07.10.737730

**Authors:** Pauline Hiort, Astrid Weiss, Janina Krentz, Ralph T Schermuly, Harm-Jan Bogaard, Tim Conrad, Robert Szulcek, Katharina Baum

## Abstract

Pulmonary arterial hypertension (PAH) represents a heterogeneous group of disorders that involves complex molecular dysregulations, which are not fully captured by single-omics analyses. We apply our network-based multi-omics analysis framework, DrDimont, to transcriptomic, proteomic, phosphoproteomic, and kinase screening data from lung microvascular endothelial cells of PAH patients and controls.

Thereby, we extend the functionality of DrDimont to incorporate kinase–kinase interactions during the construction of condition-specific multi-omics networks. Kinase interactions are inferred from phosphorylations of screened substrates that are weighted by kinase-substrate predictions. Differential interaction scores from the network-based analysis between PAH and control uncover alterations centered on kinases, in particular top hits relating to MAPK signaling, such as MAPK13, MAP2K, or upstream IRAK1, and other MAPK/MAP2K family members. Further highly differential nodes were ACADSB, GPX7, DSE (for proteins), and AIM1, LY96, CHSY3 (for mRNAs). When prioritizing drug candidates by mapping drug targets onto the differential network, we find high scores for the drug tacrolimus (FK506) and several anti-neoplastic MAPK inhibitors (e.g., selumetinib, trametinib), as well as agents acting on general proliferation via (mitochondrial) DNA transcription (e.g., epirubicin, topotecan).

Integrating kinase activity screens into our explainable multi-omics network-based analyses reveals kinase-centered alterations and therapeutic hypotheses in PAH that complement single-layer classical differential expression analyses.

## 1 Introduction

Pulmonary arterial hypertension (PAH) is a rare vascular lung disorder where pulmonary arteries are remodeled and obstructed progressively [Ghofrani et al., 2025; Humbert et al., 2023; Mocumbi et al., 2024]. The increased blood pressure leads to higher strain on the right ventricle of the heart and can ultimately result in heart failure [Ghofrani et al., 2025; Humbert et al., 2023]. The PAH pathophysiology is associated with genetic, epigenetic, and environmental factors [Ghofrani et al., 2025], and the molecular mechanisms are highly heterogeneous [Humbert et al., 2023]. Thus, the diagnosis and treatment of PAH pose a challenge.

First investigations on the molecular basis of PAH’s pathophysiology have been started [Mocumbi et al., 2024]. Different omics data layers have been examined: transcriptomics (e.g., Singh et al. [2023]; Stearman et al. [2019]; Wittig et al. [2025]), proteomics (e.g., Abdul-Salam et al. [2010]; Rhodes et al. [2017]; Xu et al. [2019]), or phosphoproteomics and metabolomics (e.g., Xu et al. [2019]).

The majority of these studies employ classical differential expression analysis in single layers, with methods such as DESeq2 [Love et al., 2014] for transcriptomics data, for identifying molecular mechanisms of PAH [Abdul-Salam et al., 2010; Rhodes et al., 2017; Stearman et al., 2019; Xu et al., 2019]. However, given the broad realm of molecular alterations underlying PAH, an integrative multi-omics analysis could provide better insights. As exemplified by Xu et al. [2019], a combined analysis of proteomics, phosphoproteomics, and metabolomics data suggested pathways such as the antioxidant response to be less functional in PAH. When integrating bulk and single-cell transcriptomics with genome-wide association studies, Hong et al. [2024] found, e.g., ASPN via its anti-proliferative action as a putative protective factor in a gene network analysis. However, these analyses were linear comparisons based on individual genes. There are a multitude of computational methods for integrative multi-omics analysis to uncover intricate interdependencies and non-linear relations, reviewed e.g. in Baião et al. [2025]. For example, multi-omics factor analysis (MOFA; Argelaguet et al. [2018]) relies on matrix factorization to integrate omics layers, DIABLO [Singh et al., 2019] suggests discriminant analysis if a categorical outcome variable is available, enabling, e.g., identification of biomarkers. Further, network-based methods such as similarity network fusion (SNF; Wang et al. [2014]) that mainly targets the similarity of patients, or NEMO [Rappoport and Shamir, 2019] relying on similarity in node neighborhoods have been proposed. The main advantage of network-based approaches lies in their concerted consideration of effects in many related molecules and layers together, thus summarizing and amplifying signals that are weak and could not be identified in isolation by considering their interrelations. Specifically, our previously published method, DrDimont (Drug response prediction for differential multi-omics networks), follows this principle: it constructs integrative multi-omics networks and then relies on their differential analysis to identify differential regulation across omics layers, and to prioritize potential new drugs by overlaying drug targets onto the differential multi-omics network, considering changes in an entire interactive signaling network [Hiort et al., 2022].

First-line standard therapies for PAH aim for mitigating PAH symptoms, for example, via vasodilatation, to reduce disease severity, slow its progression, and extend survival [Ghofrani et al., 2025]. However, therapies that modulate and correct the underlying molecular disease mechanisms promise more sustainable treatment. As such, sotatercept was recently approved as a novel disease-modifying add-on therapy for PAH. Alterations in kinases, such as enhanced activity of cyclin-dependent kinases (CDKs), have been implicated with PAH [Weiss et al., 2019, 2021], and in general, kinases have been under investigation as potential disease-modifying drug targets for a while now [Ghofrani et al., 2025; Weiss et al., 2021]. There is a need for more effective, causative treatments for PAH. One previously tested approach, that was used successfully in cancer therapy, is the use of kinase inhibitors, aka anti-neoplastic drugs, which target abnormal cell signaling pathways that are associated with the pathogenesis of PAH [Weiss et al., 2021]. Several deregulated protein kinases, enzymes that are crucial in cell signaling by phosphorylating specific target molecules, are linked to the hyper-proliferative microvascular endothelial cell phenotype in PAH. However, kinases are highly structurally conserved and homologous, making it challenging to target specific kinases with pharmacological inhibitors. Furthermore, neither the etiology behind the modified kinase activity nor the kinase activity patterns specific to microvessels are known in PAH. Thus, many of the systemically acting pan-kinase inhibitors evaluated in pre-clinical and clinical studies for PAH have had limited success and often suffer from severe adverse events [Weiss et al., 2021], and sometimes even induce PAH.

Here, multi-omics data from human lung microvascular endothelial cells (MVEC) of PAH and control patients is analyzed in an integrative manner. We consider transcriptomics, proteomics, phosphoproteomics, and data from a kinase-substrate screening assay. To leverage all layers together and emphasize weak but concerted signals, we employ DrDimont for a network-based multi-omics differential analysis. First, we aim to detect mRNAs, proteins, or kinases that are highly differential between PAH and control based on multi-omics evidence. An advantage of DrDimont is its ability to prioritize potential drugs, as we have shown for breast cancer previously [Hiort et al., 2022]. Second, using our method, we scrutinize the potential of approved and experimental PAH medications and investigate which anti-neoplastic drugs might be suitable for addressing the PAH molecular phenotype. Finally, we exemplify how DrDimont allows us to retrieve the molecular basis of its differential predictions and drug prioritizations (explainability).

## 2 Data and methods

### 2.1 Molecular data acquisition and preprocessing

Our dataset consists of two patient groups, PAH and control, and four omics layers comprising transcriptomics, proteomics, phosphoproteomics, and kinase substrate profiling.

#### Biological samples

Biological samples were derived and cultured as described previously in Wittig et al. [2025] and Szulcek et al. [2020], including ethics approval and written informed consent from all participants. In brief, primary human pulmonary microvascular endothelial cells (MVECs) were isolated from lungs of patients with pulmonary arterial hypertension (PAH) and from non-diseased control lung tissue. Cells were cultured under standardized and fully supplemented endothelial cell culture conditions prior to multi-omics analyses. The patient cohorts differed between omics layers, with partial overlap between datasets; samples used for proteomic and phosphoproteomic analyses were fully matched.

#### Transcriptomics

Bulk RNA-sequencing was performed as previously described [Szulcek et al., 2020]. In detail, total RNA was purified using the MagMAX-96 for Microarrays Total RNA Isolation Kit (Ambion) according to the manufacturer’s instructions, including genomic DNA removal using MagMAXTurboDNase buffer and TURBO DNase. Poly(A)+ mRNA was isolated using the Dynabeads mRNA Purification Kit (Invitrogen). Strand-specific RNA sequencing (RNA-seq) libraries were prepared using the ScriptSeq mRNA-Seq Library Preparation Kit (Epicentre). Libraries were amplified by 12 cycles of PCR and sequenced on an Illumina HiSeq2000 platform using multiplexed single-end 33 bp reads. Raw sequence data (BCL files) were converted to FASTQ format using Illumina CASAVA 1.8.2. Reads were demultiplexed based on barcode sequences, quality-controlled using FastQC, and mapped to the human reference genome (hg38) using ArrayStudio. Reads mapping to sense-strand exons were summarized at the gene level. FPKM values were used for downstream analyses with DrDimont.

#### Proteomics and phospho-proteomics

Proteins and post-translational modifications were measured using a TMT 11-plex global proteomics approach with SDC-based cell lysis (including phosphatase inhibitors and benzonase), tryptic digestion, and high-pH fractionation. Phosphopeptide enrichment was performed using the IMAC/BRAVO platform. Samples were analyzed on a Thermo Exploris mass spectrometer using a 110 min LC gradient. Raw data were processed with MaxQuant. A pooled reference channel consisting of a mixture of all samples was included in each TMT plex to enable inter-plex normalization. TMT reporter ion intensities of proteins and phosphosites were log_2_-transformed, normalized to the reference channel (log_2_(sample) - log_2_(reference)), and subsequently equal-loading normalized using a column-wise median z-score normalization. Overall, coverage was very good, with *>*9,000 quantified proteins and *>*14,000 quantified phosphosites.

#### Kinase substrate screen

Kinase substrate screening was performed using the PamStation®12 platform (Pamgene, ‘s-Hertogenbosch, The Netherlands) as previously described by Weiss et al. [2019]. Cells were lysed in M-PER lysis buffer supplemented with protease and phosphatase inhibitors. Both PTK (phospho-tyrosine kinase) and STK (serine/threonine kinase) PamChip® arrays were used. Each PTK chip contained 196 substrate peptides, and each STK chip 144 substrate peptides immobilized on a porous membrane. Kinase-mediated peptide phosphorylation was detected using phospho-site-specific fluorescent antibodies. Image acquisition and instrument control were performed using Evolve software (Pamgene), whereas image processing, quality control, signal quantification, and normalization were carried out using BioNavigator6 software (Pamgene). Individual phosphopeptide spot intensities were quantified densitometrically, quality control and exposure-time scaling were applied to exclude false-positive signals, and signals from multiple measurement cycles were integrated into a single value per peptide and sample. After background correction, peptide phosphorylation intensities were log_2_-transformed and used for downstream analyses.

#### Further processing of the molecular data

We filtered all proteins by computing the interquartile range (IQR) for each protein measurement over all samples and removing the bottom 40% of the proteins with the lowest IQR. Before IQR-based filtering, all identified proteins without a corresponding gene name were removed (13 of the total 8,974 protein measurements). We used the measurements with the highest “Intensity” in case of duplicate protein measurements. This yielded 5,377 proteins with unique gene names. In addition, we retained all measured proteins from a set of 1,065 genes of interest, including drug targets and disease-related genes, regardless of their IQR (see section 2.5). After filtering the protein data to 5,400 unique genes, the phospho-proteomics data was filtered to the respective genes/proteins in the reduced proteomics data set (yielding 7,714 measurements of 2,112 genes). In the transcriptomics data, genes were removed if they had less than two non-zero measurements per patient group (resulting in 15,551 genes of a total of 30,125). Next, the transcriptomics data was filtered to the genes in the filtered proteomics dataset together with all measured entities of the genes of interest. Thus, yielding 5,446 genes in total for the analysis (see Supplemental Table S1 for numbers of samples and measurements per layer). We removed two measurements of an artificial peptide sequence from the kinase substrate screening data, thus retaining 253 of 255 measurements.

### 2.2 Classical differential expression analysis

We conducted a classical differential expression analysis of the transcriptomics, proteomics, and phosphoproteomics data. For differential expression of the transcriptomics data, we used the R package DESeq2 (version 1.46.0) [Love et al., 2014]. We applied DESeq2 with default parameters on the read counts of our transcriptomics dataset. We employed the R package DEP (version 1.28.0) [Zhang et al., 2018] for differential analysis of the proteomics and the phosphoproteomics dataset. Here, we also used default parameters. Classical differential expression results in the volcano plots show (uncorrected) p-values; a gene is considered significantly differentially expressed if its Benjamini-Hochberg corrected p-values <0.05 (multiple-testing correction was applied separately per layer).

### 2.3 Settings for DrDimont

DrDimont, an R package, was used for a network-based multi-omics differential analysis of our dataset [Hiort et al., 2022]. DrDimont builds an integrated differential molecular network based on correlations of measured molecules in multiple omics layers (see Fig. 1A for an overview of all steps in DrDimont). The correlations serve as edge weights to build a network with molecules as nodes for each layer.

**Figure 1:**
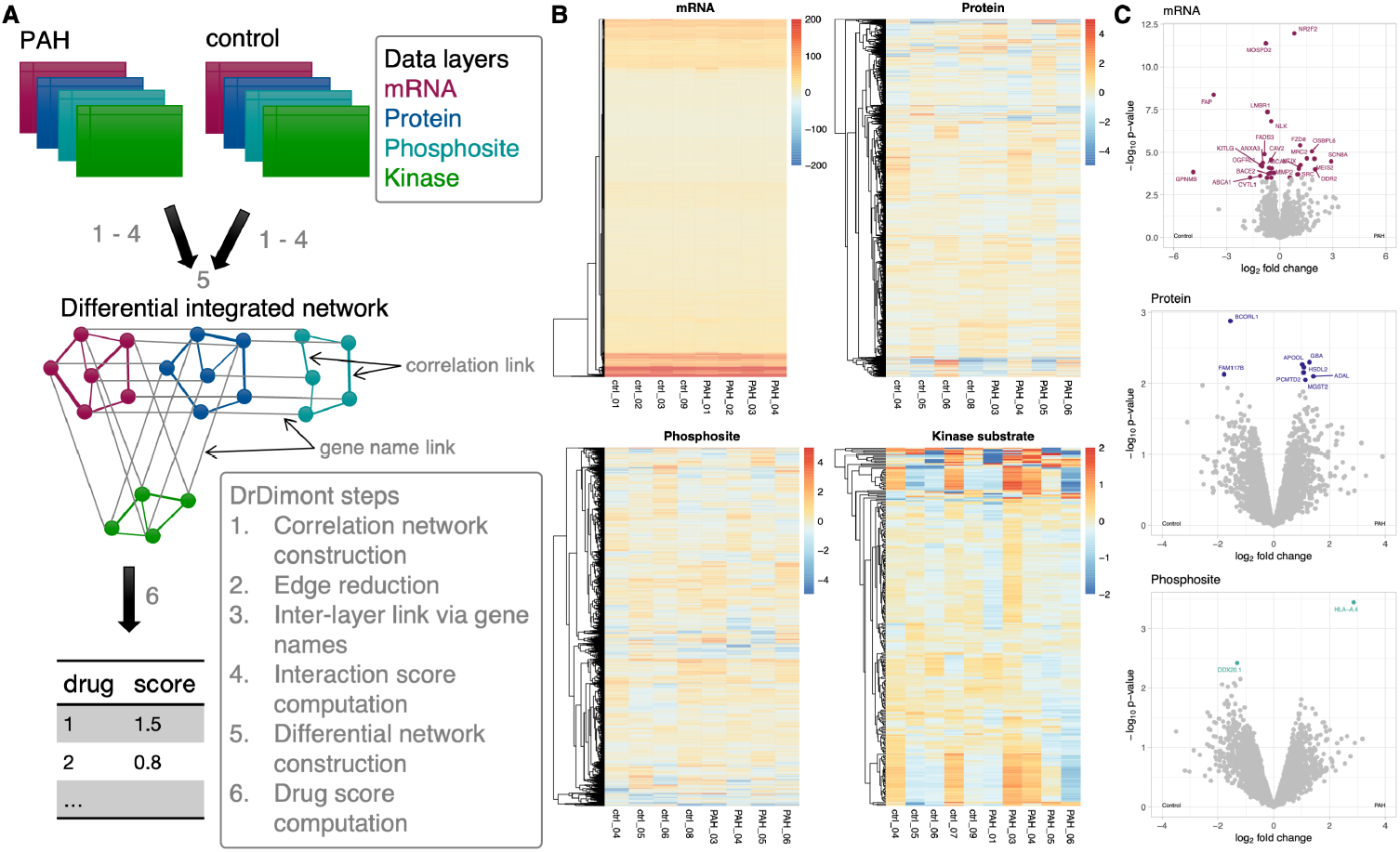
DrDimont approach and classical data overview. (A) Overview of the approach with DrDimont [Hiort et al., 2022]. For each condition (PAH and control), a condition-specific network is created by integrating the four data layers (DrDimont steps 1-4). In the networks, a node corresponds to a measured molecular entity. Drug scores are derived from the differential network based on information on the molecular targets of the drugs. (B) Clustered heatmap of the measured mRNA abundances and protein, phosphosite, and kinase substrate intensities for the control and PAH samples after filtering. Except for the kinase layer, no strong differences between conditions are observed, highlighting the need to include kinase data. (C) Volcano plots of classical differential expression analysis of filtered mRNA, protein, and phosphosite datasets. There are few significant differences (highlighted in color; significance based on Benjamini-Hochberg corrected p-values *<*0.05), but no systematic effect on specific pathways.

For our dataset, correlation matrices of the transcriptomics, proteomics, and phosphoproteomics layers were computed for the PAH and the control group, each using Pearson correlation (step 1). For the kinase layer, we built a kinase-kinase co-activity network and integrated it into the analysis after the correlation network creation with DrDimont (see section 2.4 for details).

Next, each of the correlation networks was reduced to reflect only highly correlated edges (step 2). For the edge reduction in all four layers, we used the default “pickHardThreshold” method with custom thresholds for each layer and group (see Supplemental Table S2 for thresholds). The thresholding method determines a cut-off for the correlations where the resulting reduced network reaches a given R^2^ or mean number of edges threshold. In step 3, the different omics layers were combined into a multi-omics network as a basis for a differential analysis that spans all omics layers. In our analysis, the single omics network layers were connected based on the gene identity as follows: mRNA was connected to protein, protein to phosphosite, protein to kinase, and mRNA to kinase.

The integrated interaction score computation for replacing network edges is an intermediate step in DrDimont, which adds information about alternative paths into the edge weights (step 4). Thereby, edge information reflects the local network neighborhood, which also includes other omics layers. The integrated interaction score for the edge between nodes *u* and *v, iis*_*uv*_, is defined as follows:

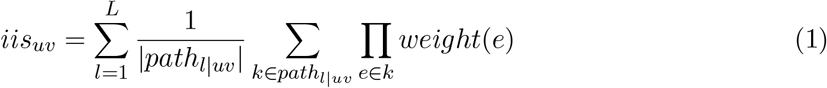

*L* is the maximum path length, *path*_*l*|*uv*_ the set of alternative paths between nodes *u* and *v* of length *l*, |*path*_*l*|*uv*_ | the number of alternative paths, and *weight*(*e*) the edge weight of edge *e*. Here, the integrated interaction scores were computed with DrDimont’s default maximum path length of three.

Next, the differential network was constructed by subtracting edge weights of the integrated interaction score network of the control group from the network of the PAH group (step 5), i.e., shown in Eq. 2.

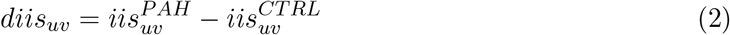

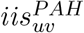 is the integrated interaction score between nodes *u* and *v* in the PAH network, and 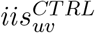 the integrated interaction score in the control network.

Finally, the differential network was used to compute differential scores for single nodes (see Eq. 3), e.g., a protein, or the differential drug score (see Eq. 4) of DrDimont (step 6).

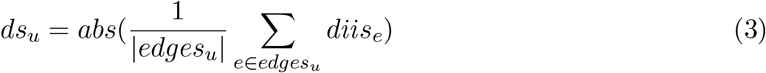

*edges*_*u*_ are all incident edges of node *u*, |*edges*_*u*_| is the number of edges incident to *u, diis*_*e*_ is the differential interaction score of edge e, and *abs*() the absolute value.

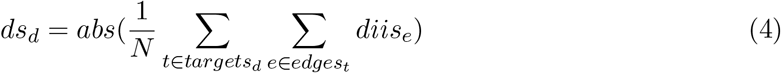

*N* is the number of all incident edges of all target nodes of drug *d, targets*_*d*_ are all targets of *d, edges*_*t*_ are all incident edges of drug target *t, diis*_*e*_ is the differential interaction score of edge *e*, and *abs*() is the absolute value.

### 2.4 Kinase layer construction

As described above, the kinase layer in our datasets consists of indirect measurements of kinase activity via a kinase substrate screen. The kinase substrate measurements are not directly comparable, making a direct comparison in DrDimont’s framework not ideal. Therefore, we decided to adapt step 1 in DrDimont to build a kinase-kinase correlation network based on kinase predictions for the given substrates and substrate-substrate correlations.

#### Kinase prediction

We constructed the kinase layer using a targeted kinase substrate screen. Possible kinases active in the samples were predicted using the Python package The Kinase Library (version 1.1.1) [Johnson et al., 2023; Yaron-Barir et al., 2024]. The Kinase Library predicts kinase-substrate interactions for 403 kinases based on binding site sequences (motifs). The 253 given peptide sequences of the substrate screen were centered around the phosphorylation site closest to the center of the respective peptide for The Kinase Library input. If present, surrounding phosphorylation sites were considered during kinase prediction using the phospho-priming feature. For other parameters, the default was used. One of the peptides from the screen could not be used for prediction by The Kinase Library because it contained an invalid amino acid. For each kinase substrate, we extracted the top 15 and 8 predicted serine/threonine and tyrosine kinases, respectively, based on the percentiles output of The Kinase Library prediction. The number of top predictions considered is in line with specifications described by The Kinase Library authors [Johnson et al., 2023; Yaron-Barir et al., 2024].

#### Kinase-kinase network construction

A matrix of interactions (e.g., correlation) between all kinases for the PAH patient group, as well as control samples, is required as input for the DrDimont framework (more details above). To overcome the limitations of the kinase activity assay, we first computed correlations of the substrate intensity measurements for each pair of substrates for each patient group. To build a kinase-kinase correlation matrix for DrDimont, we then used the correlations between their kinase substrates and the percentiles of the top Kinase Library predictions as weights (see Eq. 5). We then averaged the sum of the weighted correlations between all substrates of both kinases.

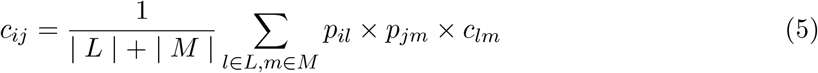

Here, *i* and *j* are predicted kinases, *L* is the set of substrates for which kinase *i* is predicted and *M* the set of substrates for which kinase *j* is predicted, *p*_*il*_ is the percentile of the prediction of kinase *i* for substrate *l, p*_*jm*_ is the percentile of the prediction of kinase *j* for substrate *m* and *c*_*lm*_ is the correlation between substrates *l* and *m*. After building the kinase-kinase correlation-like matrix, the matrix was further analyzed and integrated with the other data layers in the DrDimont framework.

### 2.5 PAH, cardiovascular, and cancer drugs and disease-related genes

#### PAH drugs

We collected a list of PAH-related drugs and their targets from literature [Ghofrani et al., 2025; Humbert et al., 2023] and based on expert knowledge. Our list consists of 16 drugs considered the standard of care in treating PAH, including sotatercept as a newly approved add-on therapy to standard-of-care treatment, and 18 experimental drugs under investigation for PAH. The latter contains drugs examined or to be examined within clinical trials for PAH from https://clinicaltrials.gov/, and the top 10 clinical phase 2 trial tyrosine kinase inhibitors (with available binding constants KD) from CHEMBL targeting kinases relevant for PAH (ERK1, ERK2, ERK5, JNK1, JNK3, p38) but at the same time not targeting those kinases relevant for proper function of healthy tissue (VEGFR1, VEGFR2, FLT3) according to the kinase activity screen. Additionally, drug targets for PAH-related drugs were compiled from CHEMBL [Zdrazil et al., 2024] and DrugBank [Wishart et al., 2017] (websites accessed on May 12, 2025). In total, we collected 142 drug-target interactions involving 34 drugs and 84 unique targets (see Supplemental Table S3 for a full list of drugs and targets).

#### Cardiovascular drugs

We gathered cardiovascular drugs and their targets from CHEMBL [Zdrazil et al., 2024] and DrugBank [Wishart et al., 2017]. Cardiovascular drugs from DrugBank (v5.1.13, accessed February 28, 2025) were identified via their ATC code starting with ‘C’. We removed any drug with the ‘withdrawn’ flag. Finally, we collected drug targets with known pharmacological action from DrugBank. Additionally, we extracted drug targets for DrugBank’s cardiovascular drugs from CHEMBL via their Python API (accessed June 10, 2025). Only drugs with proteins as their targets were included. We combined drug targets from DrugBank and CHEMBL for a more comprehensive list. In the end, 252 drugs with 803 drug target interactions were extracted.

#### Cancer drugs

We extracted anti-cancer drugs and their targets from DrugBank [Wishart et al., 2017] (version 5.1.13, access: February 28, 2025) via the ATC code ‘L01’. Drug targets were collected similarly to the cardiovascular drugs. Additionally, anti-cancer drugs and their targets were extracted from CHEMBL [Zdrazil et al., 2024] via their Python API (accessed March 10, 2025). In contrast to cardiovascular drugs, we also included drugs from CHEMBL with a cancer indication still in the experimental phase. Therefore, CHEMBL drugs were filtered to include the second-level ATC code L01 or the drug indication for neoplasm (MESH ID: D009369). Drugs were only considered for this indication if the maximum clinical phase was 3 or higher. This resulted in a total of 445 anti-cancer drugs from CHEMBL. We then extracted drug targets for these drugs and combined them with the data from DrugBank, retaining 358 drugs with 1240 drug target interactions.

#### Disease and loss-of-function genes

PAH-relevant disease genes were collected via the disease search option from the STRING protein database [Szklarczyk et al., 2023] (version 12.0; website access on February 03, 2025). We combined the results from these searches: pulmonary hypertension (DOID:6432), primary pulmonary hypertension (DOID:14557), and pulmonary arterial hypertension (HP:0002092). We identified 125 proteins associated with PAH from STRING.

In addition to the disease genes from the STRING database, we gathered genes known to have loss-of-function mutations in PAH patients as defined by Welch et al. [2023] and expert opinion. Hence, we analyzed the following 13 genes: ABCC8, ACVRL1, AQP1, ATP13A3, BMPR2, CAV1, EIF2AK4, ENG, GDF2, KCNK3, SMAD9, SOX17, TBX4.

## 3 Results

To scrutinize the molecular basis of the PAH disease mechanism, molecular measurements of four data layers of pulmonary MVEC cells cultured from PAH patients and control patients were performed (see *Data and methods* for details): mRNA expression via RNA-seq, mass-spectrometry-based shotgun proteomics and phospho-proteomics, as well as a kinase substrate screen. After filtering and quality control, we acquired data for at least four samples per condition and layer (Fig. 1B, see Supplemental Table S1 for details).

### 3.1 Classical single-layer differences between PAH and control are inconclusive

Earlier studies of us [Wittig et al., 2025] and others, e.g., Singh et al. [2023], have shown that mRNA measurements can provide insights into the molecular basis of PAH-related disease mechanisms. Top differentially expressed mRNAs from our data feature genes like NR2F2, FZD8, and OSBPL6 that are overexpressed in PAH, or MOSPD2, FAP, and LMBR1, being overexpressed in control samples (marked in color in Fig 1C top; significance based on Benjamini-Hochberg corrected p-values *<*0.05). When investigating differentially expressed proteins and phosphosites as frequent additional data layers supplementing mRNA expression, fewer significances are observed, e.g., FAM117B (downregulated protein in PAH), GBA, APOOL, HSDL2 (upregulated proteins in PAH), or HLA and DDX20 (the only statistically significantly differentially phosphorylated proteins between PAH and control conditions). In fact, the lists of differentially expressed genes do not overlap between data layers. In addition, no pathway turned up as enriched for differentially expressed genes in either the mRNA, protein, or phosphosite layer. Thus, general mechanisms or pathways responsible for the disease phenotype cannot be inferred from the three classical data layers.

### 3.2 Multi-omics network-based DrDimont can be extended to kinase data

Thus, to detect more subtle, but concerted differences in the context of regulatory networks, we applied our integrative, network-based analysis tool DrDimont [Hiort et al., 2022]. DrDimont relies on comparing omics data between conditions by creating integrated, multi-omics networks from co-expression for each condition and subsequently scrutinizing the differential network between conditions (see Fig. 1A for a pipeline overview). The network-based approach naturally allows to map molecular differences to differential drug scores for drugs with known targets, providing indications on potentially suitable drugs. In contrast to multiple other methods, such as MOFA [Argelaguet et al., 2018], DrDimont does not require any matched samples between omics layers.

When performing the kinase screen in addition to transcriptomics and proteomics measurements, we observed that differences between substrate expression of PAH and control are manifold in this layer (see Fig. 1B bottom). Therefore, we extended DrDimont to incorporate kinase screening data (see Fig. 2A for an overview). In particular, the single-omics condition-specific network generation (DrDimont step 1, see Fig. 1A) requires adaptation: In contrast to the other data layers for which the abundances of entities represented as nodes are measured directly, kinase activity is assessed indirectly via intensity measurements of potential substrates. As intensity measurements of different kinase substrates are not comparable, directly inferring kinase activity from them is not advisable and might hide relevant signals.

**Figure 2:**
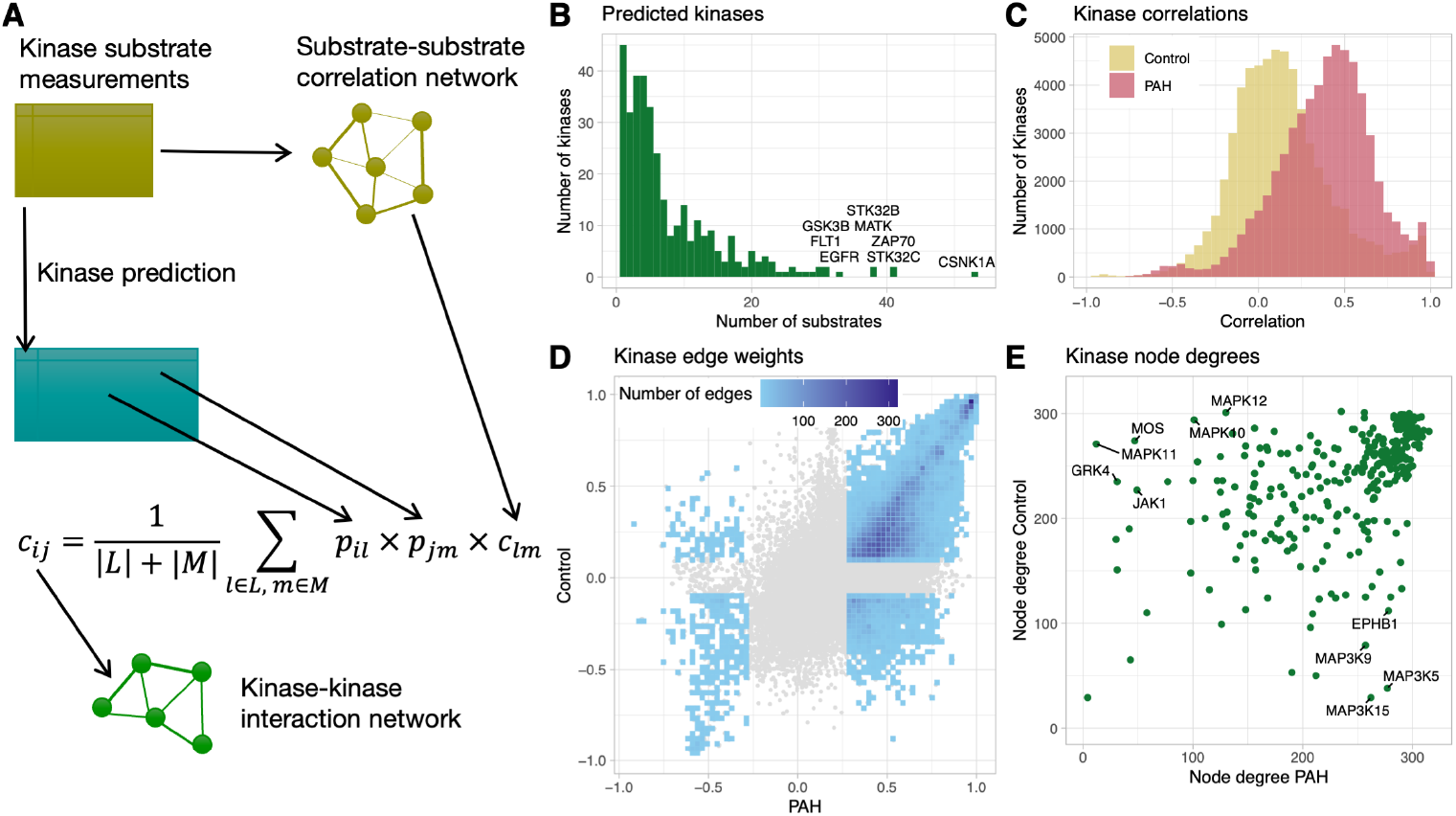
Creation and properties of the kinase interaction networks in PAH vs. control. (A) Overview of kinase-kinase interaction network construction from kinase-substrate measurements. In order to normalize kinase substrate intensity measurements, they are used to create a weighted PAH and a control substrate-substrate network based on correlations of the intensities. Percentiles *p*_*il*_ of a kinase *i* being responsible for the phosphorylation of substrate *l* are predicted with The Kinase Library [Johnson et al., 2023; Yaron-Barir et al., 2024]. The kinase-kinase network for PAH and for control is then created by averaging for each pair of kinases the edge weights of all different pairs of substrates, weighted by the kinase-substrate probabilities. (B) Distribution of the number of substrates for which a kinase is predicted among the top hits by The Kinase Library (see *Data and methods*). CSNK1A is the kinase with the most associated substrates (*>*50). (C) Edge weight distributions of the created kinase-kinase interaction networks for PAH vs. control. Overall, the PAH network displays stronger interactions between kinases. (D) Density plot directly comparing edge weights of the kinase interaction network of PAH vs. control. Edge weights show a positive correlation between conditions. In grey, low-weight edges are shown that are removed via thresholding for the subsequent analysis (see *Data and methods*). (E) Degrees of the kinase nodes in the resulting kinase-kinase interaction networks for control vs. PAH.

Therefore, we designed a three-step process to infer a condition-specific kinase-kinase interaction graph to use with DrDimont: (i) map kinases to their substrates, (ii) create a substrate co-phosphorylation graph, (iii) merge the substrate-specific kinase mapping and the substrate graph to build a kinase-kinase graph.

In detail, for the first step, we used the tool The Kinase Library [Johnson et al., 2023; Yaron-Barir et al., 2024] to predict for each measured substrate those kinases that are held responsible for the substrate’s phosphorylation (see *Data and methods* for more details). This prediction exclusively relies on (modified) sequence information of the substrates, not their measured intensities, and on systematic experimental characterizations of binding sites of kinases [Johnson et al., 2023; Yaron-Barir et al., 2024]. Among the 403 possible kinases that are supported within The Kinase Library, the majority are predicted for only a few kinase substrates from the screen (Fig. 2B). However, some kinases act promiscuously, such as CSNK1A, ZAP70, or STK32C, which are predicted for 40 or more kinase substrates.

Second, from the substrate intensity measurements, we generated weighted substrate-substrate correlation networks for PAH and for control conditions. We used the WGCNA approach [Langfelder and Horvath, 2008] in the DrDimont pipeline, as we did for the other data layers.

Third, for each kinase-kinase pair, we employed the predicted percentiles of the kinase-substrate pairs from The Kinase Library as weights and averaged over all associated substrate-substrate edge weights to infer the kinase-kinase interaction weight of the pair (see Fig. 2A and *Data and methods*). Iterating over the kinase pairs, a fully connected, weighted kinase-kinase interaction network for each condition was created. The resulting networks differ for control vs.

PAH, with PAH exhibiting overall stronger kinase-kinase interactions (Fig. 2C). Subsequently, less important, i.e., low-weight kinase-kinase interaction edges were truncated (Fig. 2D) while keeping biological network properties, following a similar procedure as for the other data layers in DrDimont.

The resulting degrees of the kinase nodes in the kinase-kinase interaction networks for control vs. PAH are overall coherent for many kinases (upper right in Fig. 2E). However, for some kinases, the degrees between PAH and control differ considerably. In particular, MAP3K5, MAP3K9, MAP3K15, and EPHB1 are decisively more connected in the PAH network (more than 300 connections in PAH vs. 100 or fewer in control), whereas kinases such as MAPK11 or GRK4 are more connected in the control network. This hints at potential alterations and thus disease relevance for PAH in the regulations of the associated pathways.

### 3.3 A differential network analysis provides insights into molecular mechanisms

For a differential multi-omics network-based analysis of the data with DrDimont, the single-layer networks for PAH and control of the transcriptomics, proteomics, and phosphosite data layers were created from the data (see *Data and methods*). Together with the kinase-kinase network, they were connected to a multi-omics network for each condition, with the proteomic layer at the center. For this, mRNA and protein nodes with matching gene identities were linked, as well as kinase and protein nodes with matching gene identities. Further, protein nodes were linked to the nodes of their corresponding phosphorylated forms in the phosphosite layer. These contributed links were incorporated as edges of weight one. We did not connect the kinase and phosphosite layer via the kinase-substrate relationship due to only a very limited number of overlapping entities. Connected networks were integrated with DrDimont using a semi-local edge weight propagation approach where alternative paths up to length three are considered. This results in edge weights of the integrated multi-omics networks, so-called interaction scores, between -3 and 3 (see *Data and methods* and [Hiort et al., 2022]), denoting strong negative or positive relationships between nodes. Note, however, that these created networks are not causal, but only phenomenologically represent the observed measurements and their statistical connections.

The characteristics of the resulting multi-omics networks for PAH and control, including their resolution by data layer, are shown in Fig. 3A. Thereby, the kinase layer is the smallest with the fewest nodes. The differential network for PAH vs. control (its properties are depicted in Fig. 3B-D) represents the differences in the integrated multi-omics networks for the two conditions, mainly based on their edge weights (see *Data and methods*, Eq. 2). Non-existent edges are thereby treated as edges with an interaction score of zero. Differential interaction scores range from -6 to 6, with a value of 6, for example, indicating a strong positive interaction for PAH, and a strong negative interaction in the control network.

**Figure 3:**
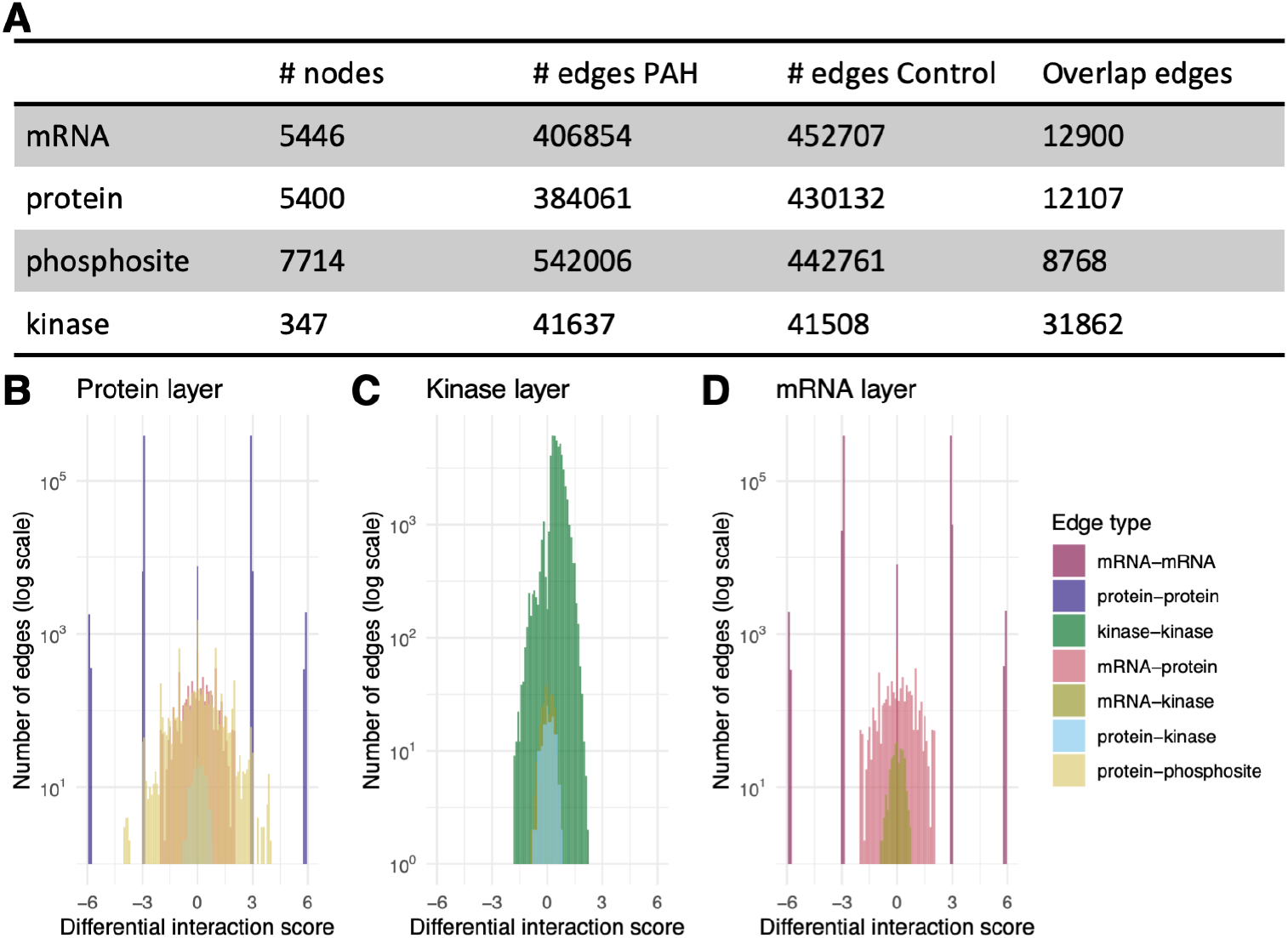
Characteristics of the differential network PAH vs. control. (A) Number of nodes from each layer. While the nodes are identical between PAH and control multi-omics integrated networks, edges are derived from the measurements and thus represent differences in their connectivities between conditions. Except for the kinase layer, there is only a small overlap between edges within the other layers. As nodes are identical, also between-layer edges fully overlap between conditions (not reported). (B-D) Histograms of the differences of the interaction scores between PAH and control, resolved by the types of nodes an edge connects (edge type). High differential scores indicate stronger connectivity in PAH than in control. Note that we do not consider interaction scores of phosphosite-phosphosite edges, as these are not of relevance for subsequent differential node predictions. See Supplemental Fig. S1 for interaction scores for each single condition.

Overall, mRNA-kinase edges (Fig. 3C, D) and protein-kinase edges (Fig. 3B, C) tend to exhibit differential interaction scores closest to zero, followed by mRNA-protein (Fig. 3B, D) and protein-phosphosite edges (Fig. 3B). Edges within mRNA and protein layers show nearly discrete distributions, with almost exclusively taking values close to -6, -3, 0, 3, 6 (Fig. 3B, D). This probably results from the strict threshold when truncating the single-omics networks, leaving exclusively edges of weights very close to -1 and 1, and topological restrictions on possible alternative paths between nodes that enforce them to include mainly edges from prior knowledge of weight one and being of maximal (or minimal) weight.

The differential network gives rise to alternative, network-based differentiality assessments for the modeled molecular entities that are derived from average differential interaction scores adjacent to the nodes in question (see *Data and methods*, Eq. 3, and Fig. 4). Nodes with particularly high differential interaction scores have particularly different connections and weights when comparing the PAH and the control network. Note that the differential interaction score analysis provides a complementary perspective to differential abundance analysis: When comparing, e.g., the list of top differential proteins according to DrDimont (Fig. 4A) with those from the classical differential expression (Fig. 1C middle), there is no protein overlap (see also Supplemental Fig. S3).

**Figure 4:**
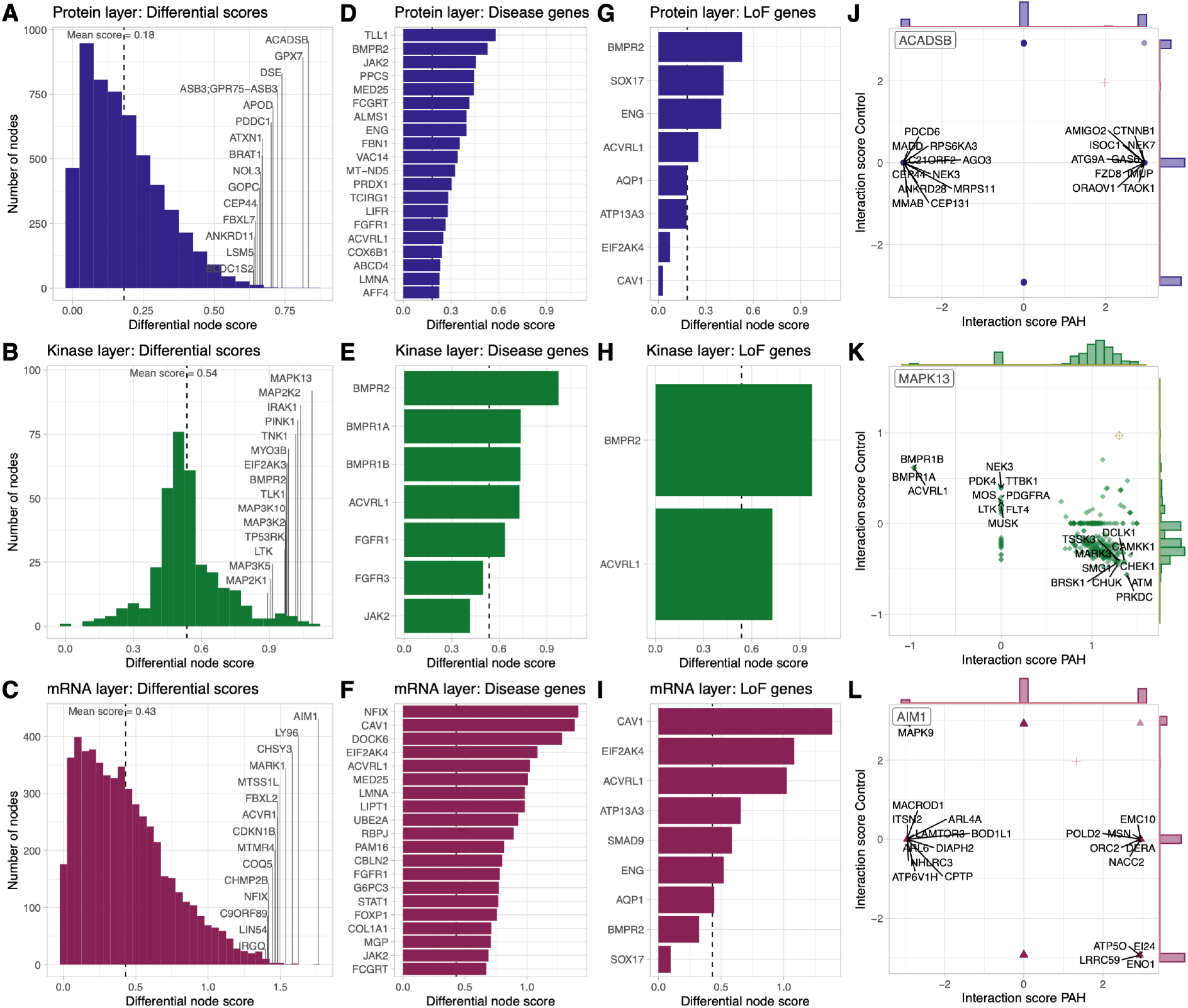
Differential scores of nodes in three omics layers. A, D, G, J: protein, B, E, H, K: kinase, C, F, I, L: mRNA layer. Differential scores for a node were computed by averaging all weights of its adjacent edges in the differential network (see Eq. 3). For each data layer, the full histogram of nodes (A-C), their restriction on PAH-related genes (as defined in STRING; D-F; only top 20 genes in D and F), or known loss-of-function (LoF) genes in PAH (G-I). The dashed line shows the mean differential node score for the respective layer. Explanations of the differential score for a node rely on mapping the relevant edges back to their weights in the control vs. PAH network and allow for diagnosing the origin of strong differentiality (shown for the most differential node of each layer; J-L).

Overall, between PAH and control, we find ACADSB, GPX7, and DSE as top differentially connected proteins (Fig. 4A), MAPK13, MAPK2K, IRAK1 as top differentially connected kinases (Fig. 4B), and AIM1, LY96, CHSY3 as top differentially connected mRNAs (Fig. 4C). Of note, four more kinases related to MAPK signaling (MAP3K10, MAP3K2, MAP3K5 and MAP2K1) appear among the top 15 differentially connected kinase nodes. We observe the strongest differential scores in the mRNA layer.

We investigated network-based differentiality of PAH-related disease genes mapped to our molecular entities (according to the STRING database; see *Data and methods*). While often not among the top-scorers, at least 35 percent of the disease-related entities revealed above-average differential scores for each layer (32/79 proteins, 5/7 kinases, 33/95 mRNAs; top20 shown in Fig. 4D-F; Supplemental Fig. S2 for all disease genes in protein and mRNA layers). In terms of overlap between layers, the kinase node BMPR2 scores eighth most differential in the kinase layer, and second differential among the disease-related protein nodes, and the disease-relevant NFIX gene is the 12th most differential mRNA node overall. Top-scoring known loss-of-function (LoF) genes in PAH are BMPR2 and ACVRL1 across all three layers, CAV1 scores high in the mRNA layer, but lowest of all LoF genes on the protein level (Fig. 4G-I).

The DrDimont approach can explain these results by investigating the root cause of differentiality (Fig. 4J-L). For ACADSB, for example, a strong rewiring between PAH and control conditions drives network-based differentiality: strong negative connections of ACADSB with proteins like PDCD6, NEK3, and positive connections to FZD8, NEK7, and CTNNB1 arise for PAH compared to control, and some relationships are lost (interaction score of zero for PAH, gene names not shown, Fig. 4J). For the MAPK13 kinase node, differentiality arises, among others, from its positive connections to the kinases BMPR1B, ACVRL1 in control shifting to negative ones in PAH, and slightly negative connections to some kinases, such as ATM and PRKDC, in control are strongly positive for PAH (Fig. 4K). For the AIM1 mRNA node, of note, a strong positive connection to MAPK9 in control is strongly negative in PAH, and strong negative interactions to ATP5O, EI24, LRRC59, ENO1 in control are strongly positive in PAH (Fig. 4L).

### 3.4 Differential drug scores from DrDimont allow prioritizing novel therapies

We had shown previously that our multi-omics network-based differential analysis with DrDimont can help prioritize drugs between breast cancer subtypes [Hiort et al., 2022]. Thus, we hypothesized that a similar analysis on the PAH data could also reveal drugs with a particularly high potential for addressing PAH-relevant pathways and thus treatment potential.

Therefore, we extended our network-based differential analysis to drugs. For each drug, we retrieved its known drug targets from CHEMBL and DrugBank (see *Data and methods*) and investigated their role in the differential network. We computed a drug’s differential score by averaging its targets’ differential edge weights (see Eq. 4). Thus, drugs providing a very high network-based differential score act within regions of the molecular network that are especially different in control vs. PAH. We distinguished between assuming drug targets stem from the protein layer (Fig. 5 top row), the kinase layer (Fig. 5 middle row), or the mRNA layer (Fig. 5 bottom row).

**Figure 5:**
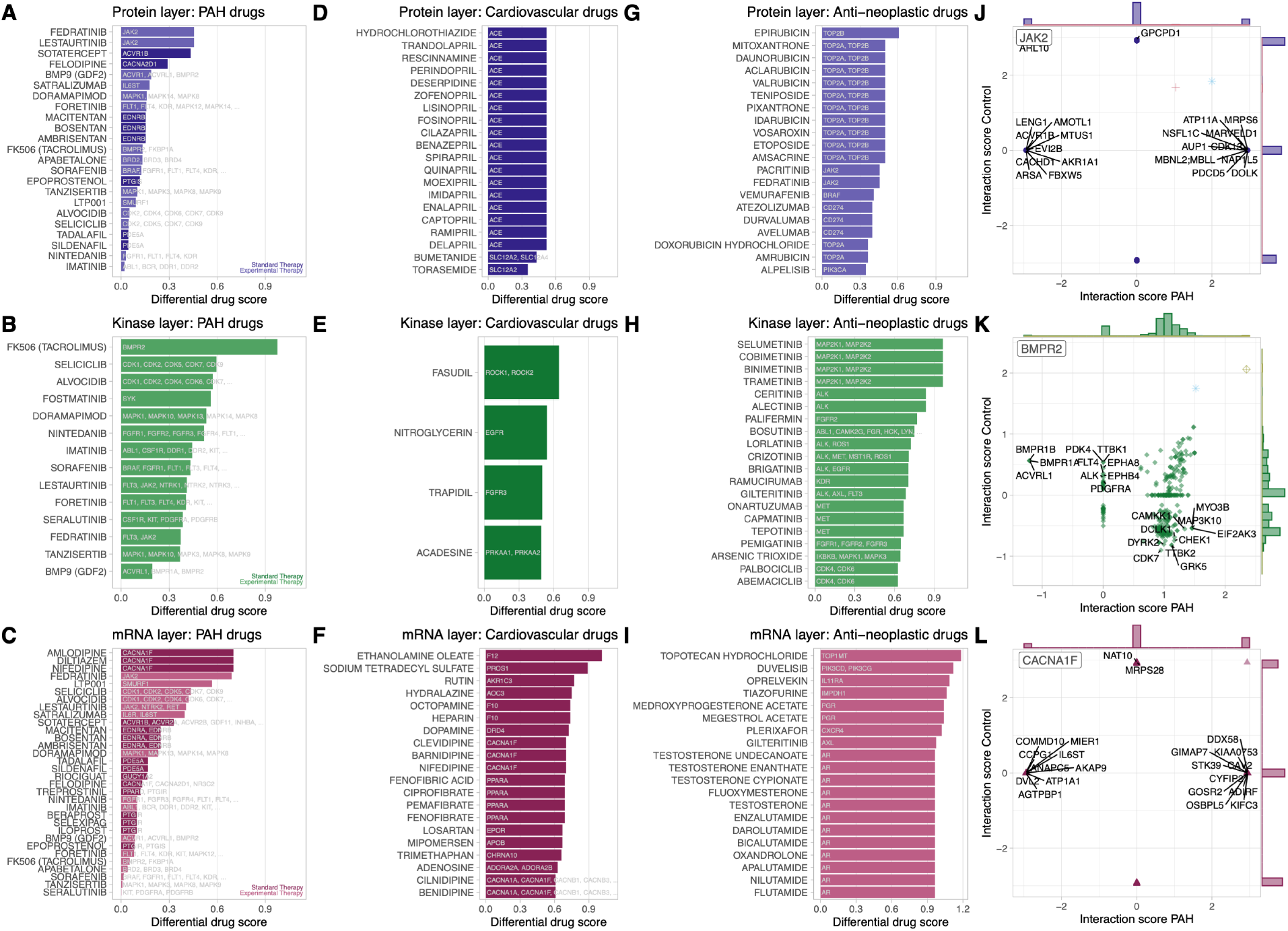
Network-based differential drug scores based on nodes in the protein layer (top row, blue), kinase layer (middle row, green), and mRNA layer (bottom row, red). We report differential scores for PAH drugs (A-C; standard therapy darker, experimental light), the top 20 cardiovascular drugs (D-F), and the top 20 anti-neoplastic drugs (G-I). For the most differential PAH drugs, the underlying differences of their drug targets in the molecular networks between PAH and control are resolved (J-L).

First, we focused on PAH drugs, both those being employed for standard therapy as well as those under consideration experimentally (see Supplemental Table S3; Fig. 5A-C). Based on the kinase and mRNA layer, overall higher drug scores than for the protein layer are observed. The drug with the highest differential score is the experimental therapy drug FK506, due to its indirect targeting of the strongly differential BMPR2 kinase node (Fig. 5B and Fig. 4B, Spiekerkoetter et al. [2013]). Particularly, network connections of BMPR2 to other kinases differ between control and PAH, such as BMPR1A, BMPR1B, ACVRL1 (strongly negative in PAH), or EIF2AK, GRK5, CHEK1 (strongly positive in PAH) (Fig. 5K). Drugs scoring second and third based on the kinase layer, seliciclib and alvocidib, are targeting CDK genes (Fig. 5B). Standard therapy drugs targeting CACNA1F - amilodipine, diltiazem, nifedipine - receive strong differential scores when evaluated on the mRNA layer (Fig. 5C). The experimental drug fedratinib that targets JAK2 reaches high differential scores in the mRNA and protein layer (Fig. 5A, C). The underlying altered connections reside mainly within the mRNA or protein layer (Fig. 5J). Interestingly, the drug lestaurtinib shows the same high drug score as fedratinib in the protein layer, but a reduced score in the mRNA layer due to further, less differential targets NTRK2, RET (Fig. 5A, C). Some standard PAH therapy drugs, such as tadalafil, sildenafil, and epoprostenol, show notably low differential drug scores. This suggests that these drugs rather act less specifically towards the PAH phenotype, but may rely on more common pathways not altered compared to healthy control, or on a systemic rather than molecular level. The same applies to the experimental drugs nintedanib and imatinib that have been described for PAH therapy before.

Second, we assessed network-based differential drug scores for cardiovascular drugs that might be commonly applied for PAH treatment. For the protein level, drugs with ACE as a drug target score highest (Fig. 5D). Fasudil, targeting ROCK1 and ROCK2, is the drug with the highest differential score in the kinase layer (Fig. 5E). When considering mRNA nodes, ethanolamine oleate and sodium tetradecyl sulfate score highly, acting via F12 or PROS1 gene, respectively (Fig. 5F).

We can use our network-based differential analysis to prioritize potential new therapies. For this, our analysis builds on PAH having been reported to share similarities with a cancer phenotype [Mocumbi et al., 2024; Weiss et al., 2021], and experimental therapies have suggested anti-neoplastic drugs such as imatinib [Ghofrani et al., 2025]. Therefore, we screened drug differential scores of all anti-neoplastic drugs reported in CHEMBL and DrugBank in each of the three data layers (see top 20 drugs according to differential score in Fig. 5G-I). The top-scoring agents are epirubicin (targeting TOP2B), selumetinib, cobimetinib, binimetinib, trametinib (targeting MAP2K1, MAP2K2), and topotecan hydrochloride (targeting TOP1MT).

## 4 Discussion

We applied our network-based approach, DrDimont, for a multi-omics integrative analysis to compare MVECs of PAH patients and controls. Therefore, we extended the framework to explicitly include a kinase-kinase network inferred from substrate screening data, and combined it with data from transcriptomic, proteomic, and phosphosite measurements.

With DrDimont’s differential network analysis that integrates statistical relationships within and across data layers, we found differentially connected entities that were not top hits according to classical single-layer differential expression analysis. For example, ACADSB, which acts in fatty acid metabolism, or GPX7, as an important player in cellular response to oxidative stress, emerge as highly differential at the protein-network level. Thus, classical differential analysis and network-based differential interaction scores are complementary: single-layer expression analyses detect individual molecular changes, whereas differential interaction scores point to possible changes of functional relationships that underlie the PAH phenotype. The kinase layer proves particularly informative for characterizing PAH, with overall higher connectivity in the PAH-related kinase-kinase interaction network. Further, MAPK13, MAPK2K, and IRAK1 (acting upstream of MAPK signaling) are the top three differentially connected kinases, and four additional MAPK signaling family member genes appear among the top 15 differential kinase nodes in our analysis. This observation suggests modulation of MAPK cascade wiring in PAH endothelial cells, which may underlie some phenotypic similarities between PAH and cancer-like proliferative states, as has been reported previously [Mocumbi et al., 2024; Weiss et al., 2021]. However, of note, also the integrative multi-omics differential investigation by DrDimont did not reveal concerted pathway-level changes for PAH vs. control, possibly due to the limited number of samples or high molecular heterogeneity among PAH patients [Humbert et al., 2023].

By mapping known drug targets onto DrDimont’s multi-omics differential network, we obtain drug-level scores that prioritize agents whose targets lie in strongly differential network regions. Applying this procedure to screen cancer drugs highlighted several candidates with potential relevance to PAH. A group of MAPK pathway inhibitors (selumetinib, cobimetinib, binimetinib, trametinib) scored highly, reflecting the possible implication of MAPK in the PAH molecular phenotype. Drugs inhibiting (mitochondrial) DNA transcription more generally, such as epirubicin (TOP1M as target) and topotecan (TOP2B as target), also ranked highly, suggesting a potential relevance of targeting proliferative signaling for PAH treatment. Interestingly, low differential scores of several standard PAH therapies (e.g., sildenafil, tadalafil, epoprostenol) may reflect that their modes of symptomatic or systemic action do not align with the cellular and molecular differences captured here. However, as we derive insights from primary cell data, even though sampled directly from patients, the in-vitro-in-vivo gap prevails at least partially, and translating our suggestions requires rigorous experimental validation, in particular, to account for toxicity and systemic effects.

Our approach carries several limitations: First, the sample size is small, which reduces statistical power and increases vulnerability to sampling variability. This issue is emphasized by PAH being a highly heterogeneous disease. As our analysis treats all PAH patients as a single group, patient-specific network changes will remain hidden. Second, DrDimont’s multiomics integration is phenomenological as network edges represent statistical associations and their propagations rather than causal relationships. Therefore, inferred differentiality should be interpreted as hypotheses of altered coordination rather than mechanistic proof. Third, the chosen connections between data layers within DrDimont are conservative: we linked layers only via direct gene or protein identity, and did not incorporate additional prior knowledge, such as from protein-protein interactions or gene regulatory networks [Müller-Dott et al., 2023; Szklarczyk et al., 2023]. While the latter could increase sensitivity to subtle, but concerted changes in pathways, it also introduces possible erroneous network information and risks biasing results toward known biology. Fourth, the drug prioritization depends on known drug-target annotations and on those targets being represented within our measurements, and it excludes the assessment of drugs without any known targets.

Overall, our approach illustrates how network-based analysis can reveal mechanisms and therapeutic hypotheses that are complementary and provide added benefit to classical single-layer differential expression analyses. Together, the data and analyses reveal kinases, and in particular the MAPK-family signaling, as highly disease-relevant in PAH, and accordingly prioritize MAPK pathway inhibitors, among others, as putative novel therapeutic options. Studies with larger sample sizes will be required to more robustly infer common PAH disease mechanisms, and more personalized insights will be possible upon the adaptation of methods like DrDimont to act on subgroup or single-sample level.

## 5 Data availability

Transcriptomics data is available from the Supplementary Material in [Wittig et al., 2025]. Further experimental data and code are available on reasonable request.

## 6 Acknowledgements

We thank Thea Steuerwald for assistance with the DrDimont analysis. Furthermore, we thank Andrea Volkamer and Philipp Mertins.

## Supplement

**Table S1:** Sample sizes, numbers of measured molecules of the four measured data layers for the pulmonary arterial hypertension (PAH) and control group (Ctrl). The measured molecules of the raw data, as well as the filtered datasets, are given. For the kinase layer, screened kinase substrates, i.e., phosphorylated peptides, are listed as measured entities. The phosphorylations are resolved by serine and threonine (st) and tyrosine (p) measurements.

| Data layer | PAH samples | Ctrl samples | Measured molecules (raw) | Measured molecules (filtered) |
| --- | --- | --- | --- | --- |
| mRNA | 4 | 4 | 30125 | 5446 |
| protein | 4 | 4 | 8974 | 5400 |
| phosphosite | 4 | 4 | 14358 | 7714 |
| kinase | 5 | 5 | 255 (149 p; 106 st) | 253 (148 p; 105 st) |

**Table S2:** DrDimont settings for network reduction with scale-freeness goodness-of-fit R^2^ thresholds and cut vector optimized to create networks with approximately the same size and one component in both groups. The kinase layer is reduced using the mean number of edges option in DrDimont because the R^2^ did not work well.

| Data layer | $R^2$ PAH | $R^2$ Ctrl | Cut vector PAH & Ctrl (step size: 0.001) |
| --- | --- | --- | --- |
| mRNA | 0.73 | 0.37 | 0.97 – 0.999 |
| protein | 0.1 | 0.23 | 0.95 – 0.999 |
| phosphosite | 0.005 | 0.05 | 0.97 – 0.999 |
| kinase | mean edges = 120 | mean edges = 120 | 0.1 – 0.6 |

**Figure S1:**
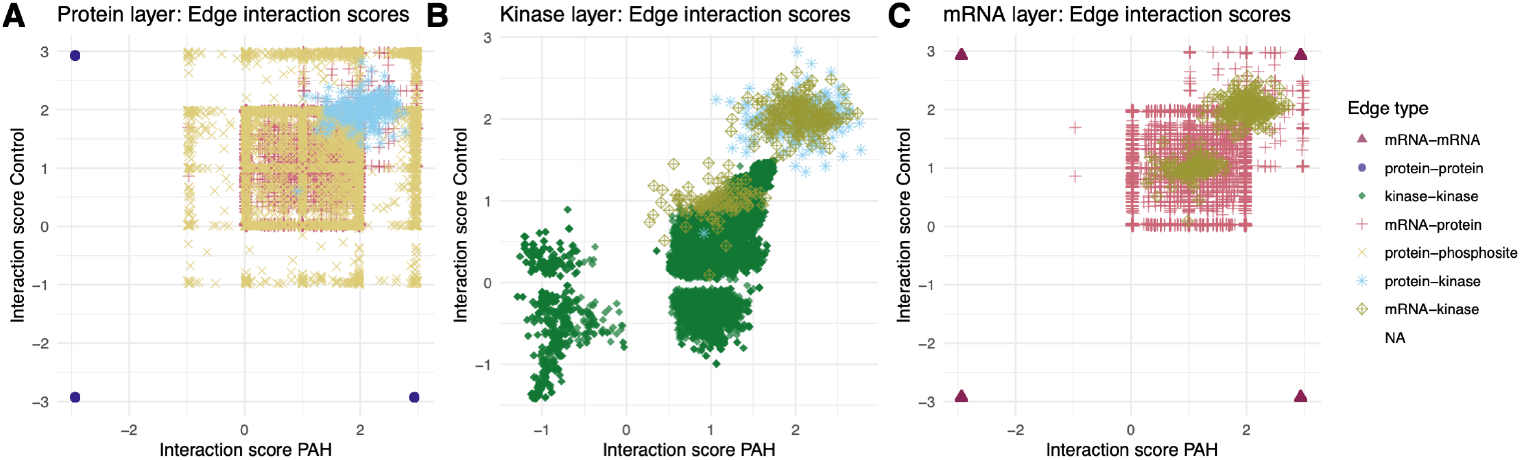
Integrated edge weight distributions of PAH and control, separated by edges adjacent to (A) protein node, (B) kinase nodes, and (C) mRNA nodes. These distributions give rise to the differential integrated edge score distributions as shown in Fig. 3.

**Figure S2:**
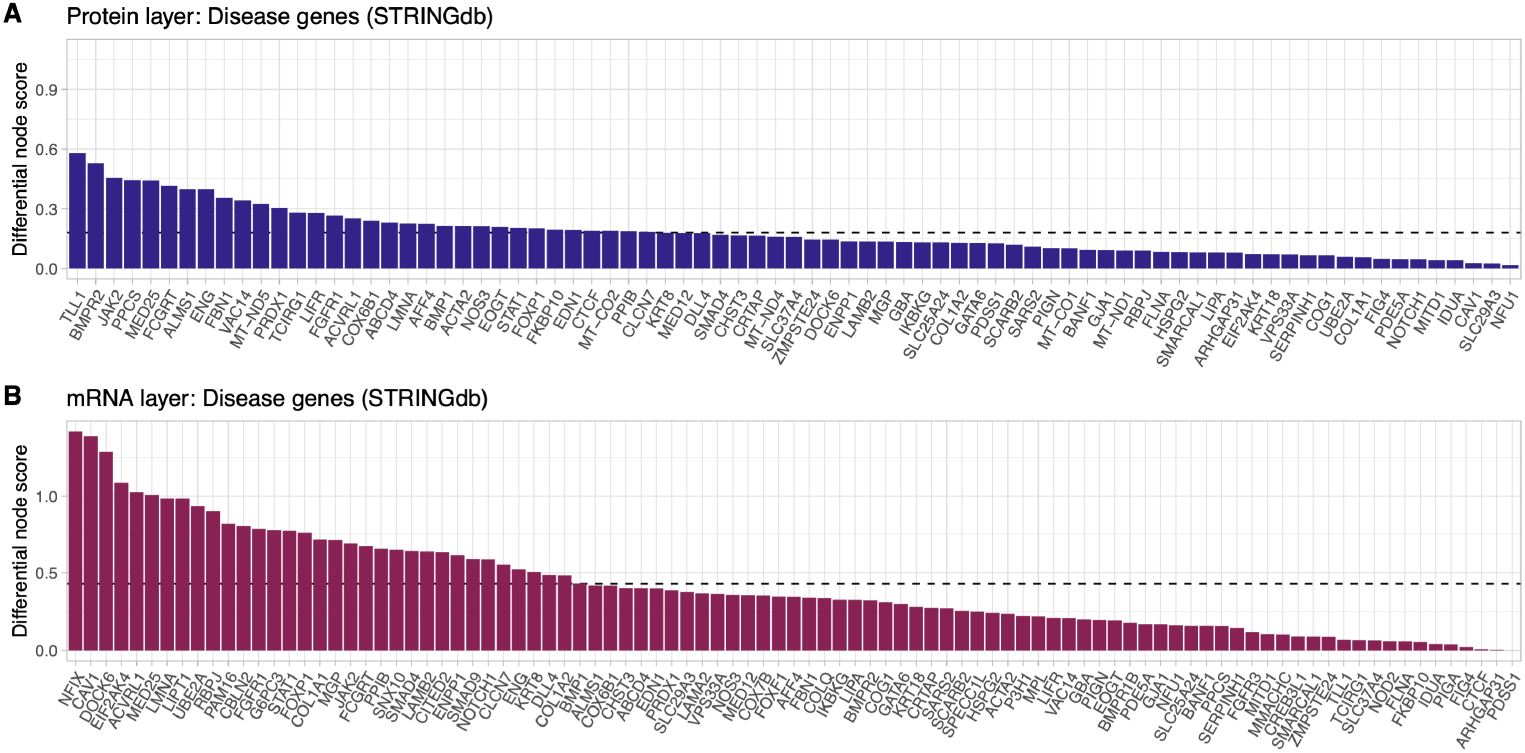
Differential scores of PAH-related disease genes from STRING in the (A) protein layer, (B) mRNA layer. The dashed line shows the mean differential score of all nodes in the respective data layer.

**Figure S3:**
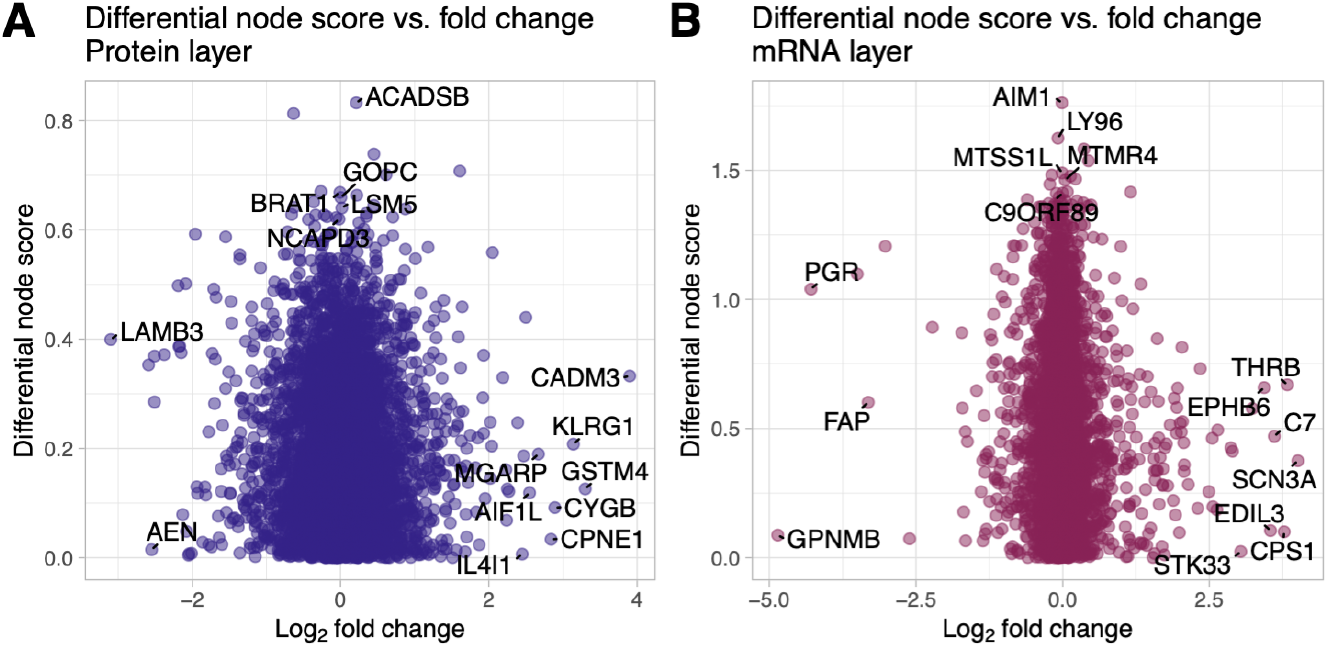
Differential scores of nodes in DrDimont’s differential network vs. the log_2_ fold change of classical differential expression analysis in the (A) protein layer, (B) mRNA layer. Shown gene names highlight genes with the highest difference between the two methods.

**Figure S4:**
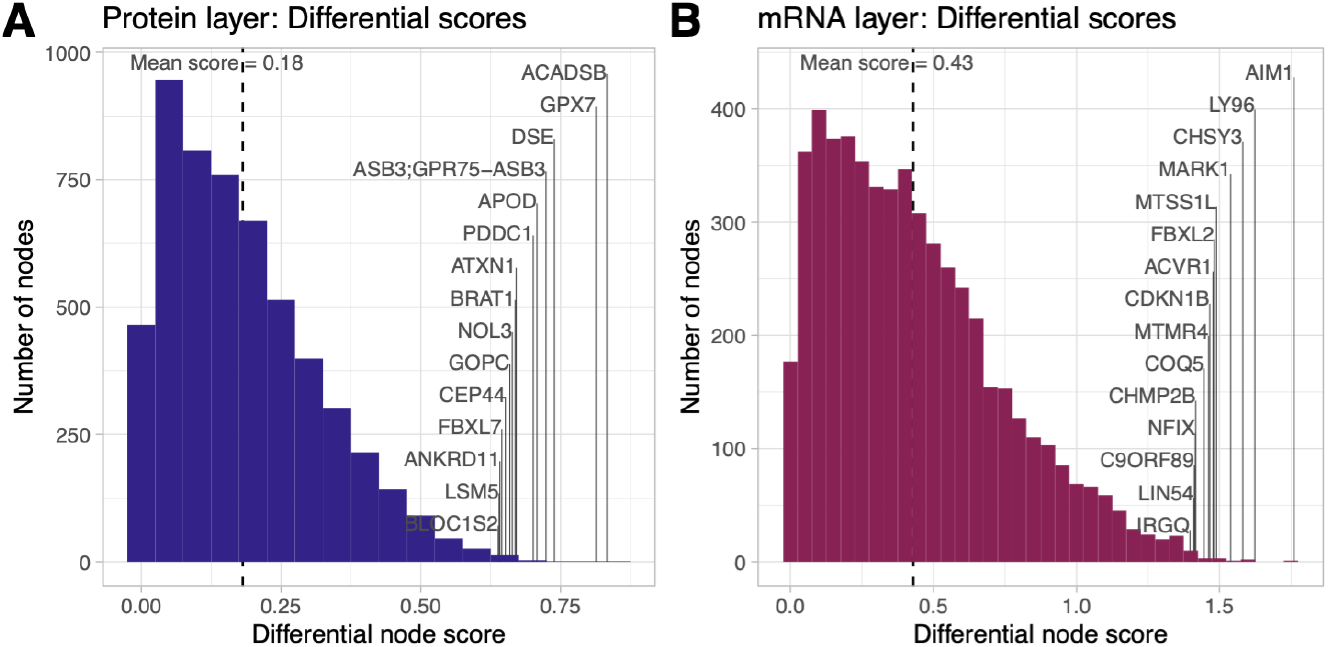
Differential scores of nodes in DrDimont’s differential network, including the mRNA, protein, and phosphoprotein layer, but not the kinase data. Distribution of differential scores in (A) protein layer and (B) mRNA layer. Shown gene names highlight nodes with the highest differential scores in each layer. Overall, in these layers, differences to the investigation, including the kinase data, are minor.

**Table S3:**
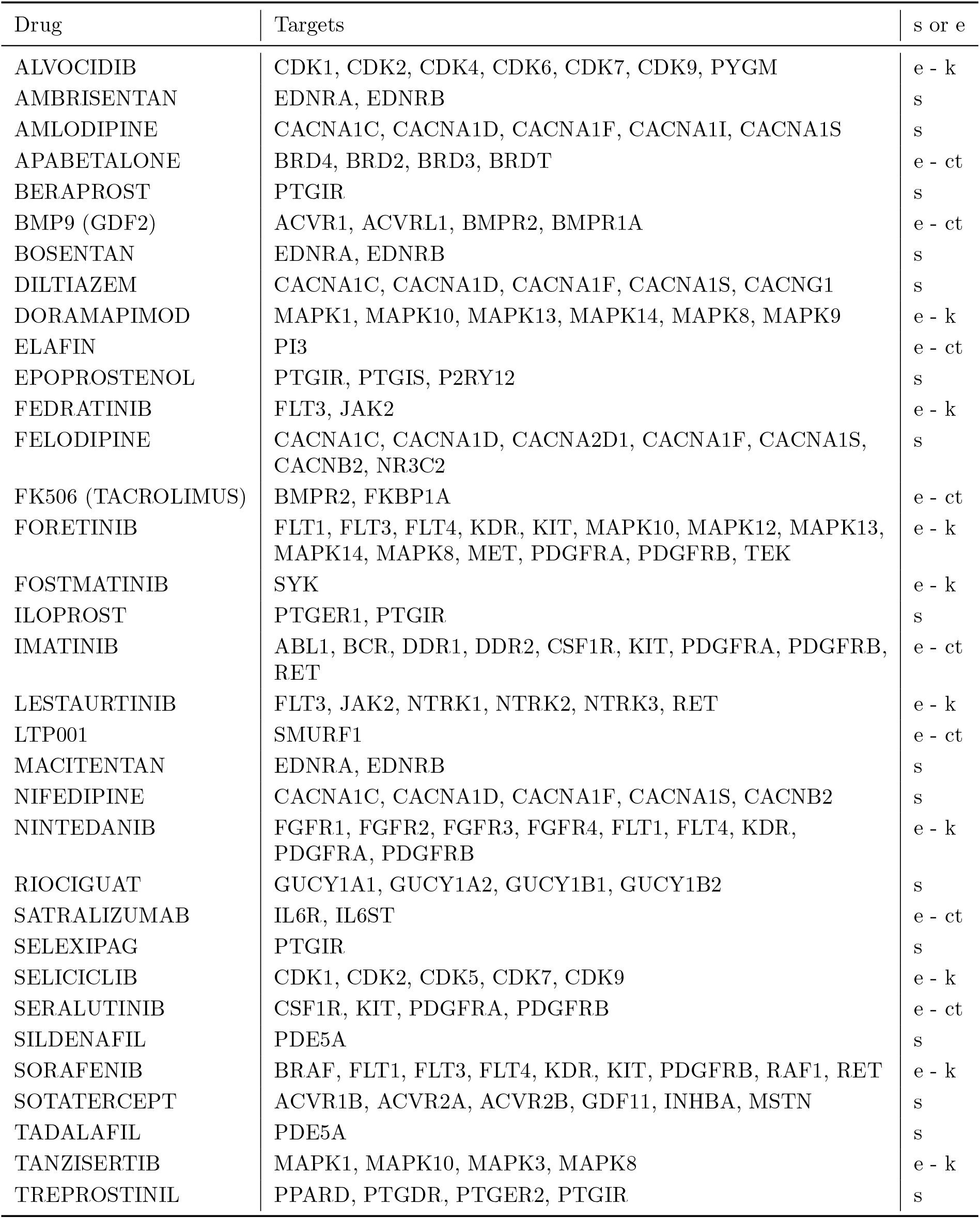
Standard and experimental drugs for PAH treatment and their targets. The list of drugs was compiled with expert help. The last column shows ‘s’ for standard therapy (from literature) and ‘e’ for experimental therapies (additionally resolving whether from clinical trials, ct, or drugs targeting kinases of interest, k).

## References

V. B. Abdul-Salam, J. Wharton, J. Cupitt, M. Berryman, R. J. Edwards, and M. R. Wilkins. Proteomic Analysis of Lung Tissues From Patients With Pulmonary Arterial Hypertension. Circulation, 122(20):2058–2067, Nov. 2010. doi:10.1161/CIRCULATIONAHA.110.972745. URL https://www.ahajournals.org/doi/10.1161/CIRCULATIONAHA.110.972745.

R. Argelaguet, B. Velten, D. Arnol, S. Dietrich, T. Zenz, J. C. Marioni, F. Buettner, W. Huber, and O. Stegle. Multi-Omics Factor Analysis—a framework for unsupervised integration of multi-omics data sets. Molecular Systems Biology, 14(6):e8124, June 2018. doi:10.15252/msb.20178124. URL https://www.embopress.org/doi/full/10.15252/msb.20178124.

A. R. Baião, Z. Cai, R. C. Poulos, P. J. Robinson, R. R. Reddel, Q. Zhong, S. Vinga, and E. Gonçalves. A technical review of multi-omics data integration methods: from classical statistical to deep generative approaches. Briefings in Bioinformatics, 26(4):bbaf355, July 2025. ISSN 1477-4054. doi:10.1093/bib/bbaf355. URL http://doi.org/10.1093/bib/bbaf355.

H.-A. Ghofrani, M. Gomberg-Maitland, L. Zhao, and F. Grimminger. Mechanisms and treatment of pulmonary arterial hypertension. Nature Reviews Cardiology, 22(2):105–120, Feb. 2025. doi:10.1038/s41569-024-01064-4. URL https://www.nature.com/articles/s41569-024-01064-4.

P. Hiort, J. Hugo, J. Zeinert, N. Müller, S. Kashyap, J. C. Rajapakse, F. Azuaje, B. Y. Renard, and K. Baum. DrDimont: explainable drug response prediction from differential analysis of multi-omics networks. Bioinformatics, 38(Supplement 2):ii113–ii119, Sept. 2022. doi:10.1093/bioinformatics/btac477. URL https://doi.org/10.1093/bioinformatics/btac477.

J. Hong, L. Medzikovic, W. Sun, B. Wong, G. Ruffenach, C. J. Rhodes, A. Brownstein, L. L. Liang, L. Aryan, M. Li, A. Vadgama, Z. Kurt, T.-H. Schwantes-An, E. A. Mickler, S. Gräf, M. Eyries, K. A. Lutz, M. W. Pauciulo, R. C. Trembath, F. Perros, D. Montani, N. W. Morrell, F. Soubrier, M. R. Wilkins, W. C. Nichols, M. A. Aldred, A. A. Desai, D.-A. Trégoüet, S. Umar, R. Saggar, R. Channick, R. M. Tuder, M. W. Geraci, R. S. Stearman, X. Yang, and M. Eghbali. Integrative Multiomics in the Lung Reveals a Protective Role of Asporin in Pulmonary Arterial Hypertension. Circulation, 150(16):1268–1287, Oct. 2024. doi:10.1161/CIRCULATIONAHA.124.069864. URL https://www.ahajournals.org/doi/10.1161/CIRCULATIONAHA.124.069864.

M. Humbert, O. Sitbon, C. Guignabert, L. Savale, A. Boucly, M. Gallant-Dewavrin, V. McLaughlin, M. M. Hoeper, and J. Weatherald. Treatment of pulmonary arterial hypertension: recent progress and a look to the future. The Lancet Respiratory Medicine, 11 (9):804–819, Sept. 2023. doi:10.1016/S2213-2600(23)00264-3. URL https://www.thelancet.com/journals/lanres/article/PIIS2213-2600(23)00264-3/fulltext#fig1.

J. L. Johnson, T. M. Yaron, E. M. Huntsman, A. Kerelsky, J. Song, A. Regev, T.-Y. Lin, K. Liberatore, D. M. Cizin, B. M. Cohen, N. Vasan, Y. Ma, K. Krismer, J. T. Robles, B. van de Kooij, A. E. van Vlimmeren, N. Andrée-Busch, N. F. Käufer, M. V. Dorovkov, A. G. Ryazanov, Y. Takagi, E. R. Kastenhuber, M. D. Goncalves, B. D. Hopkins, O. Elemento, D. J. Taatjes, A. Maucuer, A. Yamashita, A. Degterev, M. Uduman, J. Lu, S. D. Landry, B. Zhang, I. Cossentino, R. Linding, J. Blenis, P. V. Hornbeck, B. E. Turk, M. B. Yaffe, and L. C. Cantley. An atlas of substrate specificities for the human serine/threonine kinome. Nature, 613(7945):759–766, Jan. 2023. doi:10.1038/s41586-022-05575-3. URL https://www.nature.com/articles/s41586-022-05575-3.

P. Langfelder and S. Horvath. WGCNA: an R package for weighted correlation network analysis. BMC Bioinformatics, 9(1):559, Dec. 2008. doi:10.1186/1471-2105-9-559. URL https://doi.org/10.1186/1471-2105-9-559.

M. I. Love, W. Huber, and S. Anders. Moderated estimation of fold change and dispersion for RNA-seq data with DESeq2. Genome Biology, 15(12):550, Dec. 2014. doi:10.1186/s13059-014-0550-8. URL https://doi.org/10.1186/s13059-014-0550-8.

A. Mocumbi, M. Humbert, A. Saxena, Z.-C. Jing, K. Sliwa, F. Thienemann, S. L. Archer, and S. Stewart. Pulmonary hypertension. Nature Reviews Disease Primers, 10(1):1, Jan. 2024. doi:10.1038/s41572-023-00486-7. URL https://www.nature.com/articles/s41572-023-00486-7.

S. Müller-Dott, E. Tsirvouli, M. Vazquez, R. Ramirez Flores, P. Badia-i Mompel, R. Fallegger, D. Türei, A. Lægreid, and J. Saez-Rodriguez. Expanding the coverage of regulons from high-confidence prior knowledge for accurate estimation of transcription factor activities. Nucleic Acids Research, 51(20):10934–10949, 11 2023. ISSN 0305-1048. doi:10.1093/nar/gkad841. URL https://doi.org/10.1093/nar/gkad841.

N. Rappoport and R. Shamir. NEMO: cancer subtyping by integration of partial multi-omic data. Bioinformatics, 35(18):3348–3356, Sept. 2019. ISSN 1367-4803. doi:10.1093/bioinformatics/btz058. URL https://doi.org/10.1093/bioinformatics/btz058.

C. J. Rhodes, J. Wharton, P. Ghataorhe, G. Watson, B. Girerd, L. S. Howard, J. S. R. Gibbs, R. Condliffe, C. A. Elliot, D. G. Kiely, G. Simonneau, D. Montani, O. Sitbon, H. Gall, R. T. Schermuly, H. A. Ghofrani, A. Lawrie, M. Humbert, and M. R. Wilkins. Plasma proteome analysis in patients with pulmonary arterial hypertension: an observational cohort study. The Lancet Respiratory Medicine, 5(9):717–726, Sept. 2017. doi:10.1016/S2213-2600(17)30161-3. URL https://www.thelancet.com/journals/lanres/article/PIIS2213-2600(17)30161-3/fulltext.

A. Singh, C. P. Shannon, B. Gautier, F. Rohart, M. Vacher, S. J. Tebbutt, and K.-A. Lê Cao. DIABLO: an integrative approach for identifying key molecular drivers from multi-omics assays. Bioinformatics, 35(17):3055–3062, Sept. 2019. doi:10.1093/bioinformatics/bty1054. URL https://doi.org/10.1093/bioinformatics/bty1054.

N. Singh, C. Eickhoff, A. Garcia-Agundez, P. Bertone, S. S. Paudel, D. T. Tambe, L. A. Litzky, K. Cox-Flaherty, J. R. Klinger, S. F. Monaghan, C. J. Mullin, M. Pereira, T. Walsh, M. Whittenhall, T. Stevens, E. O. Harrington, and C. E. Ventetuolo. Transcriptional profiles of pulmonary artery endothelial cells in pulmonary hypertension. Scientific Reports, 13(1):22534, Dec. 2023. doi:10.1038/s41598-023-48077-6. URL https://www.nature.com/articles/s41598-023-48077-6.

E. Spiekerkoetter, X. Tian, J. Cai, R. K. Hopper, D. Sudheendra, C. G. Li, N. El-Bizri, H. Sawada, R. Haghighat, R. Chan, L. Haghighat, V. de Jesus Perez, L. Wang, S. Reddy, M. Zhao, D. Bernstein, D. E. Solow-Cordero, P. A. Beachy, T. J. Wandless, P. ten Dijke, and M. Rabinovitch. Fk506 activates bmpr2, rescues endothelial dysfunction, and reverses pulmonary hypertension. The Journal of Clinical Investigation, 123(8):3600–3613, 7 2013. doi:10.1172/JCI65592. URL https://www.jci.org/articles/view/65592.

R. S. Stearman, Q. M. Bui, G. Speyer, A. Handen, A. R. Cornelius, B. B. Graham, S. Kim, E. A. Mickler, R. M. Tuder, S. Y. Chan, and M. W. Geraci. Systems Analysis of the Human Pulmonary Arterial Hypertension Lung Transcriptome. American Journal of Respiratory Cell and Molecular Biology, 60(6):637–649, June 2019. doi:10.1165/rcmb.2018-0368OC. URL https://doi.org/10.1165/rcmb.2018-0368OC.

D. Szklarczyk, R. Kirsch, M. Koutrouli, K. Nastou, F. Mehryary, R. Hachilif, A. L. Gable, T. Fang, N. Doncheva, S. Pyysalo, P. Bork, L. Jensen, and C. von Mering. The STRING database in 2023: protein–protein association networks and functional enrichment analyses for any sequenced genome of interest. Nucleic Acids Research, 51(D1):D638–D646, Jan. 2023. doi:10.1093/nar/gkac1000. URL https://doi.org/10.1093/nar/gkac1000.

R. Szulcek, G. Sanchez-Duffhues, N. Rol, X. Pan, R. Tsonaka, C. Dickhoff, L. M. Yung, X. D. Manz, K. Kurakula, S. M. Kielbasa, H. Mei, W. Timens, P. B. Yu, H.-J. Bogaard, and M.-J. Goumans. Exacerbated inflammatory signaling underlies aberrant response to BMP9 in pulmonary arterial hypertension lung endothelial cells. Angiogenesis, 23(4):699–714, Nov. 2020. ISSN 1573-7209. doi:10.1007/s10456-020-09741-x. URL https://doi.org/10.1007/s10456-020-09741-x.

B. Wang, A. M. Mezlini, F. Demir, M. Fiume, Z. Tu, M. Brudno, B. Haibe-Kains, and A. Goldenberg. Similarity network fusion for aggregating data types on a genomic scale. Nature Methods, 11(3):333–337, Mar. 2014. doi:10.1038/nmeth.2810. URL https://doi.org/10.1038/nmeth.2810.

A. Weiss, M. C. Neubauer, D. Yerabolu, B. Kojonazarov, B. C. Schlueter, L. Neubert, D. Jonigk, N. Baal, C. Ruppert, P. Dorfmuller, S. S. Pullamsetti, N. Weissmann, H.-A. Ghofrani, F. Grimminger, W. Seeger, and R. T. Schermuly. Targeting cyclin-dependent kinases for the treatment of pulmonary arterial hypertension. Nature Communications, 10(1):2204, May 2019. doi:10.1038/s41467-019-10135-x. URL https://www.nature.com/articles/s41467-019-10135-x.

A. Weiss, M. Boehm, B. Egemnazarov, F. Grimminger, S. Savai Pullamsetti, G. Kwapiszewska, and R. T. Schermuly. Kinases as potential targets for treatment of pulmonary hypertension and right ventricular dysfunction. British Journal of Pharmacology, 178(1):31–53, 2021. doi:10.1111/bph.14919. URL https://onlinelibrary.wiley.com/doi/abs/10.1111/bph.14919.

C. L. Welch, M. A. Aldred, S. Balachandar, D. Dooijes, C. A. Eichstaedt, S. Gräf, A. C. Houwel-ing, R. D. Machado, D. Pandya, M. Prapa, M. Shaukat, L. Southgate, J. Tenorio-Castano, E. P. Callejo, K. M. Day, D. Macaya, G. Maldonado-Velez, W. K. Chung, S. L. Archer, K. Auckland, E. D. Austin, R. Badagliacca, J.-A. Barberà, C. Belge, H. J. Bogaard, S. Bon-net, K. A. Boomars, O. Boucherat, M. M. Chakinala, R. Condliffe, R. L. Damico, M. Del-croix, A. A. Desai, A. Doboszynska, C. G. Elliott, M. Eyries, M. P. Escribano Subías, H. Gall, S. Ghio, A.-H. Ghofrani, E. Grünig, R. Hamid, L. Harbaum, P. M. Hassoun, A. R. Hemnes, K. Hinderhofer, L. S. Howard, M. Humbert, D. G. Kiely, D. Langleben, A. Lawrie, J. E. Loyd, S. Moledina, D. Montani, N. W. Morrell, W. C. Nichols, A. Olschewski, H. Olschewski, S. Papa, M. W. Pauciulo, S. Provencher, R. Quarck, C. J. Rhodes, L. Scelsi, W. Seeger, D. J. Stewart, A. Sweatt, E. M. Swietlik, C. Treacy, R. C. Trembath, O. Tura-Ceide, C. D. Vizza, A. Vonk Noordegraaf, M. R. Wilkins, R. T. Zamanian, and D. Zateyshchikov. Defining the clinical validity of genes reported to cause pulmonary arterial hypertension. Genetics in Medicine, 25(11):100925, Nov. 2023. ISSN 1098-3600. doi:10.1016/j.gim.2023.100925. URL https://www.sciencedirect.com/science/article/pii/S1098360023009383.

D. S. Wishart, Y. D. Feunang, A. C. Guo, E. J. Lo, A. Marcu, J. R. Grant, T. Sajed, D. Johnson, C. Li, Z. Sayeeda, N. Assempour, I. Iynkkaran, Y. Liu, A. Maciejewski, N. Gale, A. Wilson, L. Chin, R. Cummings, D. Le, A. Pon, C. Knox, and M. Wilson. DrugBank 5.0: a major update to the DrugBank database for 2018. Nucleic Acids Research, 46(D1):D1074–D1082, Nov. 2017. doi:10.1093/nar/gkx1037. URL https://doi.org/10.1093/nar/gkx1037.

C. Wittig, J. M. König, X. Pan, J. Aman, H.-J. Bogaard, P. B. Yu, W. M. Kuebler, K. Baum, and R. Szulcek. Shear stress unveils patient-specific transcriptional signatures in PAH: Towards personalized molecular diagnostics. Theranostics, 15(5):1589–1605, Jan. 2025. doi:10.7150/thno.105729. URL https://www.thno.org/v15p1589.htm.

W. Xu, S. A. A. Comhair, R. Chen, B. Hu, Y. Hou, Y. Zhou, L. A. Mavrakis, A. J. Janocha, L. Li, D. Zhang, B. B. Willard, K. Asosingh, F. Cheng, and S. C. Erzurum. Integrative proteomics and phosphoproteomics in pulmonary arterial hypertension. Scientific Reports, 9(1):18623, Dec. 2019. doi:10.1038/s41598-019-55053-6. URL https://www.nature.com/articles/s41598-019-55053-6.

T. M. Yaron-Barir, B. A. Joughin, E. M. Huntsman, A. Kerelsky, D. M. Cizin, B. M. Cohen, A. Regev, J. Song, N. Vasan, T.-Y. Lin, J. M. Orozco, C. Schoenherr, C. Sagum, M. T. Bedford, R. M. Wynn, S.-C. Tso, D. T. Chuang, L. Li, S. S.-C. Li, P. Creixell, K. Krismer, M. Takegami, H. Lee, B. Zhang, J. Lu, I. Cossentino, S. D. Landry, M. Uduman, J. Blenis, O. Elemento, M. C. Frame, P. V. Hornbeck, L. C. Cantley, B. E. Turk, M. B. Yaffe, and J. L. Johnson. The intrinsic substrate specificity of the human tyrosine kinome. Nature, 629 (8014):1174–1181, May 2024. doi:10.1038/s41586-024-07407-y. URL https://www.nature.com/articles/s41586-024-07407-y.

B. Zdrazil, E. Felix, F. Hunter, E. J. Manners, J. Blackshaw, S. Corbett, M. de Veij, H. Ioannidis, D. M. Lopez, J. Mosquera, M. Magarinos, N. Bosc, R. Arcila, T. Kizilören, A. Gaulton, A. Bento, M. Adasme, P. Monecke, G. Landrum, and A. Leach. The ChEMBL Database in 2023: a drug discovery platform spanning multiple bioactivity data types and time periods. Nucleic Acids Research, 52(D1):D1180–D1192, Jan. 2024. doi:10.1093/nar/gkad1004. URL https://doi.org/10.1093/nar/gkad1004.

X. Zhang, A. H. Smits, G. B. van Tilburg, H. Ovaa, W. Huber, and M. Vermeulen. Proteome-wide identification of ubiquitin interactions using UbIA-MS. Nature Protocols, 13(3):530–550, Mar. 2018. doi:10.1038/nprot.2017.147. URL https://www.nature.com/articles/nprot.2017.147.

